# Contemporaneity of the past in stochastic intergenerational homeostasis

**DOI:** 10.64898/2026.04.10.717806

**Authors:** Kunaal Joshi, Karl F. Ziegler, Charles S. Wright, Elizabeth M. Spiers, Jacob T. Crosser, Shaswata Roy, Rhea Gandhi, Jack Stonecipher, Samuel Eschker, Rudro R. Biswas, Srividya Iyer-Biswas

**Author notes:** These authors contributed equally to this work (Kunaal Joshi, Karl F. Ziegler).

## Abstract

What recurring patterns of behaviors are hidden in the stochastic intergenerational dynamics of individual bacterial cells in different environments? Embracing the inherently stochastic nature of the homeostasis process, first we reconceptualize homeostasis as representing a “standing pattern of variation in trajectory space” of a naturally occurring adaptive complex system (rather than the simple “set-point” of a self-regulating apparatus), then delineate two mechanistically distinct potential routes to achieving homeostasis (either elastic, reflexive, memory-free adaptation or plastic, reflective, memoryful adaptation) and show that both schemes are simultaneously utilized during multigenerational stochastic growth and division of an individual cell. From experimental data we identify an intergenerational scaling law which directly yields the exact stochastic map governing stochastic intergenerational cell size homeostasis of individual bacterial cells. Its broad applicability across bacterial species, growth conditions, and microenvironments suggests that the organizational motif representing the nature of coupling of growth to division is effectively the same in all of these scenarios, despite apparent differences in actualization through molecular circuitry. The precise parameters characterizing the intergenerational scaling law vary from condition to condition and provide early hints of two tradeoffs: precision-speed and precision-energy.

---

How do complex systems maintain key emergent “state variables” at desired target values to within specified tolerances? This question was first posed in the context of homeostasis in living systems over a century ago. Yet the precise quantitative rules governing this phenomenon have remained fiercely debated. Here we present a direct solution to the problem through the development of a conceptual framework, precise measurements, and the extraction of emergent simplicities from quantitative data.

The idea of homeostasis was instrumental to the development of the field of physiology [1]; as formulated by Bernard in 1865, stability of an organism’s “internal milieu” is not simply a feature, but indeed a prerequisite for life [2]. Early conceptualizations philosophically permitted the possibility of stochasticity in the homeostatic variable [2–4]. However, deterministic frameworks have since become pervasive, with organisms thought—in analogy to mechanical self-regulating apparatuses—to defend the setpoint values of homeostatic variables [5] against intrinsic and extrinsic fluctuations. This approach can be traced to early developments in control theory as applied to engineered inanimate systems [6]. Within this perspective, deviations from desired setpoints are taken to indicate organismal malfunction, rather than natural variability resulting from inherent stochasticity in the dynamics [7–12].

Here we reconceptualize homeostasis as representing a “standing pattern of variation in trajectory space” of a naturally occurring adaptive complex system rather than the “setpoint” of a simple self-regulating apparatus. Embracing the inherently stochastic nature of the homeostasis process, we further delineate two mechanistically distinct potential routes to achieving it. We term the first option “elastic adaptation” or reflexive homeostasis, a passive mechanism of responding to instantaneous conditions with no retention of long-term memory (such as also exemplified by Ashby’s Homeostat [13], the first technical model of homeostasis). In contrast, active, “plastic adaptation” or reflective homeostasis is achieved through strategic integration over a finite memory of the past (as in Shannon’s Maze Machine [14]). Which scheme(s) do organisms use?

We will show that even within the context of the simplest unit of life: a bacterial cell, the most basic processes: cell growth and cell division, and the homeostases of the corresponding variables: characteristic cell size in a generation and the corresponding single-cell growth rate, *both* mechanisms, plastic (memoryful) or elastic (memoryless), are simultaneously at play! In the context of bacterial cell homeostasis, while the preponderance of interest has focused on the cell size problem, the previously reported *intra*generational scaling laws contained early hints of both homeostases [15]. Briefly, the mean-rescaled cell division times from different balanced growth conditions were found to undergo a scaling collapse, indicating the emergence of a cellular unit of time, which can be calibrated by the mean single-cell growth rate in a given condition. When appropriately rescaled in its terms the dynamics of stochastic growth and division become condition independent [15–17]. Moreover the mean-rescaled cell size distributions from different times since the last division event, and from different temperatures, were found to undergo scaling collapses, also suggesting how a single dominant timescale may arise from the underlying complex molecular circuitry governing the growth of a bacterial cell in a given balanced growth condition [15, 18]. However, these *intra*generational scaling laws do not provide insight into how stochastic *inter*generational homeostasis of either cell size or the corresponding single-cell growth rate is achieved. Furthermore, the question of how deterministic inter-generational models (such as the sizer–timer–adder family of models [19]) are to be generalized to precisely capture the observed signatures of stochasticity in cell size and growth rate dynamics remains to be fully addressed (Fig. 1 f, g). Here we address this question through the development of a conceptual framework, and a synthesis of precise measurements with extraction of emergent simplicities from quantitative data. In the following sections, to quantify the dynamics of cell growth and division on inter-generational scales, we make use of the following quantities – (i) the cell size immediately following a cell division event, labeled ‘initial size’ (*a*_*i*_), (ii) the cell size immediately before division, labeled ‘final size’ (*a* _*f*_), and (iii) the exponential growth rate of cell size between division events (*k*). These quantities are depicted in Fig. 1 c. Furthermore, we denote the initial size of consecutive generations of a single cell by *a*_*n*_, where *n* is the number of generations since the starting generation (*n* = 0).

**Figure 1:**
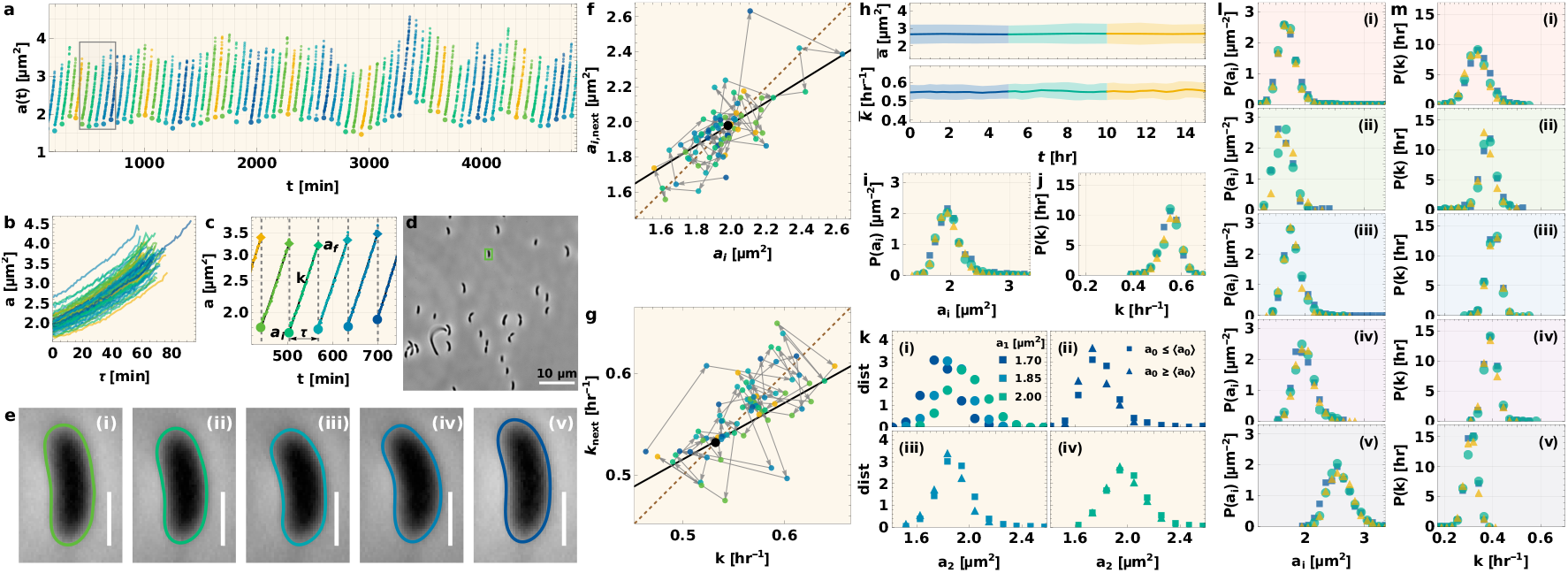
Stochastic intergenerational homeostases of the individual cell’s size and growth rate under constant growth conditions. (a) A typical individual C. crescentus stalked cell’s intergenerational growth and division trajectory obtained using our SChemostat technology [12, 15], under time-invariant balanced growth conditions. The plot shows cell size as a function of time across multiple division cycles (with separate division cycles shown in different colors). The initial sizes (cell size immediately after division) in each generation are marked by larger circles. (b) The growth trajectories for each generation are plotted as functions of cell age (time since the last division event). (c) The growth trajectory for the generations highlighted by the box in (a) plotted with a log scale for cell size. The trajectories between divisions appear linear, indicating exponential growth, with growth rate (k) given by the slope. The initial size (cell size immediately after division) and final size (cell size immediately before division) are labeled as a_i_ and a _f_ respectively, and the division time is labeled as τ. (d) Representative experimental field of view, with the cell corresponding to the trajectory in (a) highlighted. A typical experiment consists of tens of such fields of view, imaged tens of times between successive division events, with constant cell number and density for the duration of the experiment. (e) Cell boundary splines generated by our custom automated image analysis routines from phase contrast images such as those shown in (d). These boundaries are used to compute cell areas such as those shown in (a) with 𝒪 (1%) measurement precision, as detailed in [12, 15]. The different panels correspond to the initial sizes of the generations of the corresponding color in (c). Scale bar = 1µm. (f) Contrast between predictions from deterministic homeostatic models (the sizer–timer–adder family of models) and the inherently stochastic nature of intergenerational cell size dynamics. The deterministic models predict a stepwise descent towards the mean (central black point) bounded by the growth curve (black line, corresponding to the average value of next generation’s initial size (a_i,next_) given current generation’s initial size a_i_) and the division curve (dashed brown line, corresponding to a_i,next_ = a_i_). The actual cell’s trajectory behaves differently as shown. (g) Analogous plot for the single cell growth rate (k) as shown in (f) for cell size. (h) The mean cell size (upper panel) and growth rate (lower panel) along with standard deviation (shaded region) remain constant in time under constant growth conditions (complex media at 30° C). The distributions of pooled (i) initial cell size and (j) growth rate are plotted for cell generations drawn from the three time windows shown in (h), using corresponding colors. These distributions are invariant, establishing that stochastic intergenerational homeostasis has been achieved and is sustained. (k) Evidence that the intergenerational cell size dynamics are Markovian. (i) The distributions of the subsequent generation’s initial size (a_2_) given the current generation’s initial size (a_1_) are plotted for three different values of a_1_. Point colors indicate distinct values of a_1_. (ii-iv) For each value of a_1_ in (i), the conditional distribution of the subsequent generation’s initial size (a_2_) given the current generation’s initial size (a_1_) is further disambiguated according to whether the previous generation’s initial size (a_0_) is less than (square) or greater than (triangle) the population mean initial size. The distributions overlap irrespective of whether a_0_ is less than or greater than the population mean, demonstrating the Markovian property. (l) and (m) show the distributions corresponding to (i) and (j) for the following growth conditions – (ii) complex media post switch from minimal media at 30° C, (iii) dilute complex media at 30° C, (iv) minimal media at 30° C, (v) minimal media post switch from complex media at 30° C, and (vi) minimal media post switch from complex media at 25° C.

## Results

### Direct evidence of stochastic intergenerational homeostasis of individual bacterial cell sizes and cell size growth rates

Surprisingly, direct evidence for stochastic intergenerational cell-size homeostasis is not readily available in published works [19–29]. Here, we present individual-cell multigenerational data (Fig. 1 a–e) from six different balanced growth conditions obtained as described in the supplementary information on experimental methods: (i) cells in balanced growth in complex (undefined) media at 30° C; (ii) cells eventually in balanced growth in complex media at 30° C after an initial multigenerational period of growth in minimal media; (iii) cells in balanced growth in dilute complex media at 30° C; (iv) cells in balanced growth in minimal (defined) media at 30° C; (v) cells eventually in balanced growth in minimal media at 30° C after an initial multigenerational period of growth in complex media; and (vi) cells eventually in balanced growth in minimal media at 25° C after an initial multigenerational period of growth in complex media. By experimentally confirming that the distribution of cell sizes at birth (“initial size distribution”) and the single-cell size growth rate distribution are time-invariant (Fig. 1 i, j, l, m) we provide direct evidence of stochastic intergenerational homeostasis of both quantities in individual cells.

### Stochastic cell size homeostasis is maintained through elastic, reflexive, memoryfree or Markovian intergenerational dynamics

When a process is memory-free (history-independent) or Markovian, future excursions of the system are determined solely by the present value of the stochastic variable, irrespective of history [30]. Under balanced growth conditions, intergenerational cell size homeostasis is maintained through reflexive, elastic adaptation (see Fig. 1 k). Therefore, the basic equation governing the multigenerational dynamics of initial cell sizes (Mathematical Derivations section 2 and [31]) is:

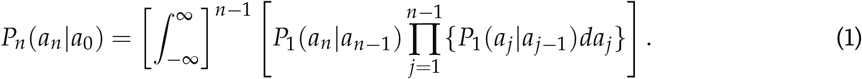

In this formulation, stable homeostasis is achieved if *P*_*n*_(*a*_*n*_|*a*_0_) converges to the same wellbehaved distribution for all values of initial cell sizes, *a*_0_, after a sufficient number of generations (*n* ≫ 1) have elapsed. This is indeed true for our cell size data in all six conditions (Fig. 2 e).

**Figure 2:**
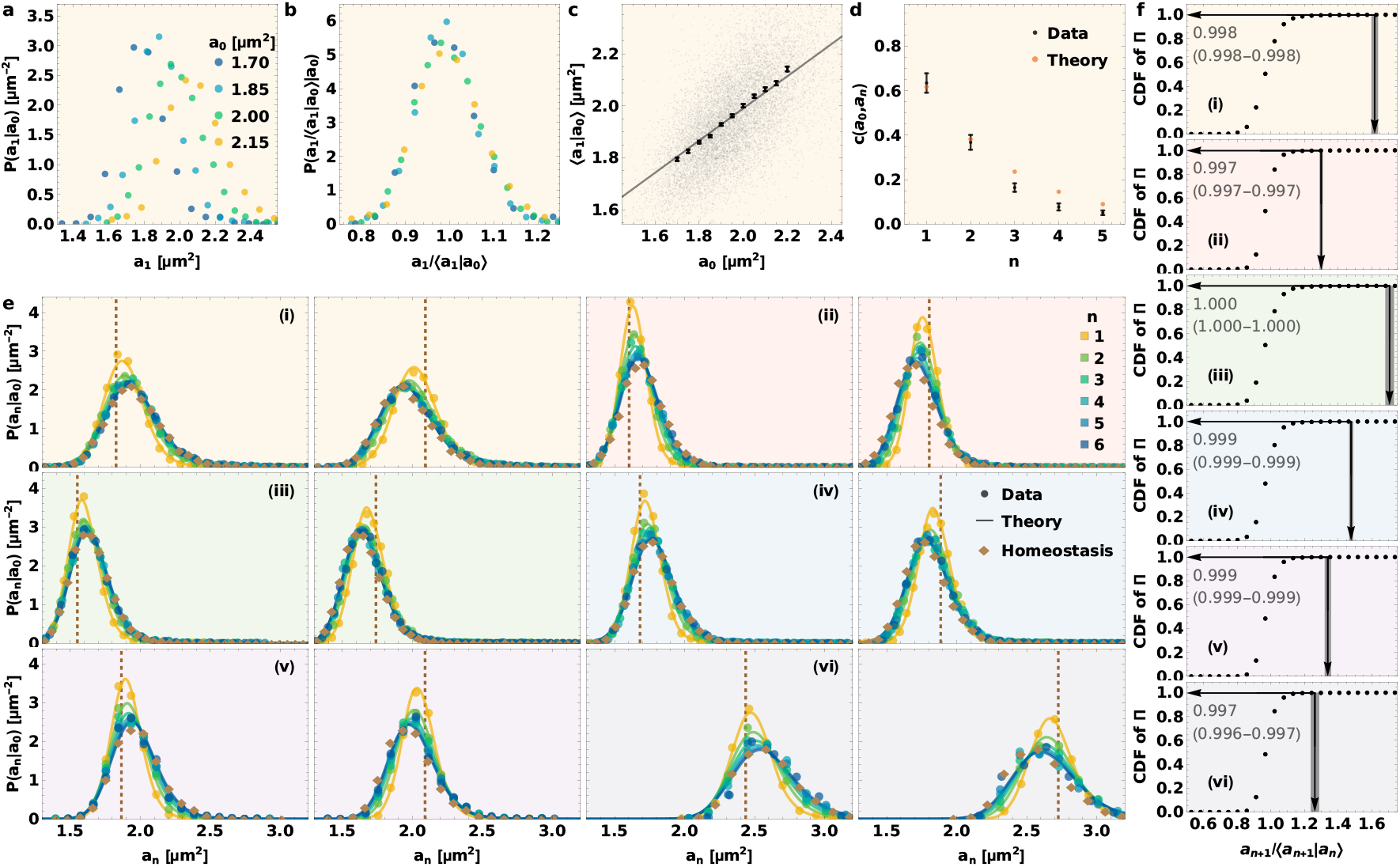
The intergenerational scaling law governing stochastic cell size homeostasis through the elastic, reflexive, memory-free scheme. (a) The distribution of the subsequent generation’s initial size, a_1_, pooled by distinct values of the current generation’s initial size, a_0_, is plotted for different values of a_0_ (different colors). (b) The distributions in (a), rescaled by their corresponding mean values, collapse to a distribution invariant of a_0_. (c) The subsequent generation’s initial size is plotted as a function of current generation’s initial size (background scatter), along with the binned means with error bars showing standard error, and the linear fit. This calibration curve is used to restore the full conditional subsequent generation’s initial size distributions conditioned on the present generation’s initial size (a) from the corresponding mean-rescaled distribution (b). (d) Coefficient of correlation between the initial sizes in the initial (zeroth) generation and after n consecutive generations for different n values. The correlations predicted by our theoretical framework (orange) closely match those measured from data (black). (e) The intergenerational evolution of the distribution of initial cell sizes, a_n_, starting from a given initial size in the starting generation (location of dashed line). n denotes the generation number, with the starting generation labeled by n = 0. Points correspond to experimental data and curves to fitting-free predictions from our theoretical intergenerational framework, which takes in as inputs the directly measured mean-rescaled distribution (b) and calibration curve (c). (i-vi) show these distributions for different growth conditions corresponding to the growth conditions shown in Fig. 1 with the corresponding background color. For each growth condition, the initial size in the starting generation is the first quartile for the panel on the left, and the third quartile for the panel on the right. (f) The cumulative distribution function (CDF) of the mean rescaled distribution (b). The value of 1/α for is marked by an arrow on the x axis (while the shaded region shows the standard error), which agrees with our theoretically derived necessary and sufficient condition for homeostasis: the upper limit of the support of the mean rescaled distribution should be less than or equal to the corresponding 1/α. Here, α is the slope of the linear fit of mean subsequent generation’s mean rescaled initial size (a_1_), conditioned on the current generation’s mean rescaled initial size (a_0_), plotted in (c). The arrow on the y axis shows the fraction of data points less than 1/α, along with the standard error. (i-vi) show the different growth conditions corresponding to (e).

### Plastic, reflective, memoryful or non-Markovian homeostasis of individual-cell growth rates

In contrast to cell size dynamics, the intergenerational temporal evolution of the single-cell growth rate in constant growth conditions is memoryful or non-Markovian. The history dependence can be revealed by considering the families of conditional probability distributions: *P*(*k*_*n*+1_ |*k*_*n*_, *k*_*n*− 1_, …), where *k*_*n*_ denotes the value of the growth rate in generation *n* and the probability that the growth rate in the (*n* + 1)^th^ generation is *k*_*n*+1_, given that the rates in the previous generations have values (*k*_*n*_, *k*_*n* −1_, …), is denoted by *P*(*k*_*n*+1_ |*k*_*n*_, *k*_*n*−1_, …). For tractability, we consider the sequence of conditional probabilities *P*(*k*_*n*+1_ |*k*_*n*_, *k*_0_), whose *k*_0_-dependence characterizes non-Markovianity due to the *n*^th^ past generation. These are shown in Fig. 3 e for generational indices *n* = 2 to *n* = 41, where *k*_*n*_ and *k*_0_ have been discretized to binary variables that are either above or below the median, 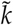, of the steady state distribution. By inspection, the difference between 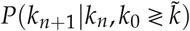, namely, the difference between the solid and dashed curves of the same color in Fig. 3 e, persists for tens of generations.

**Figure 3:**
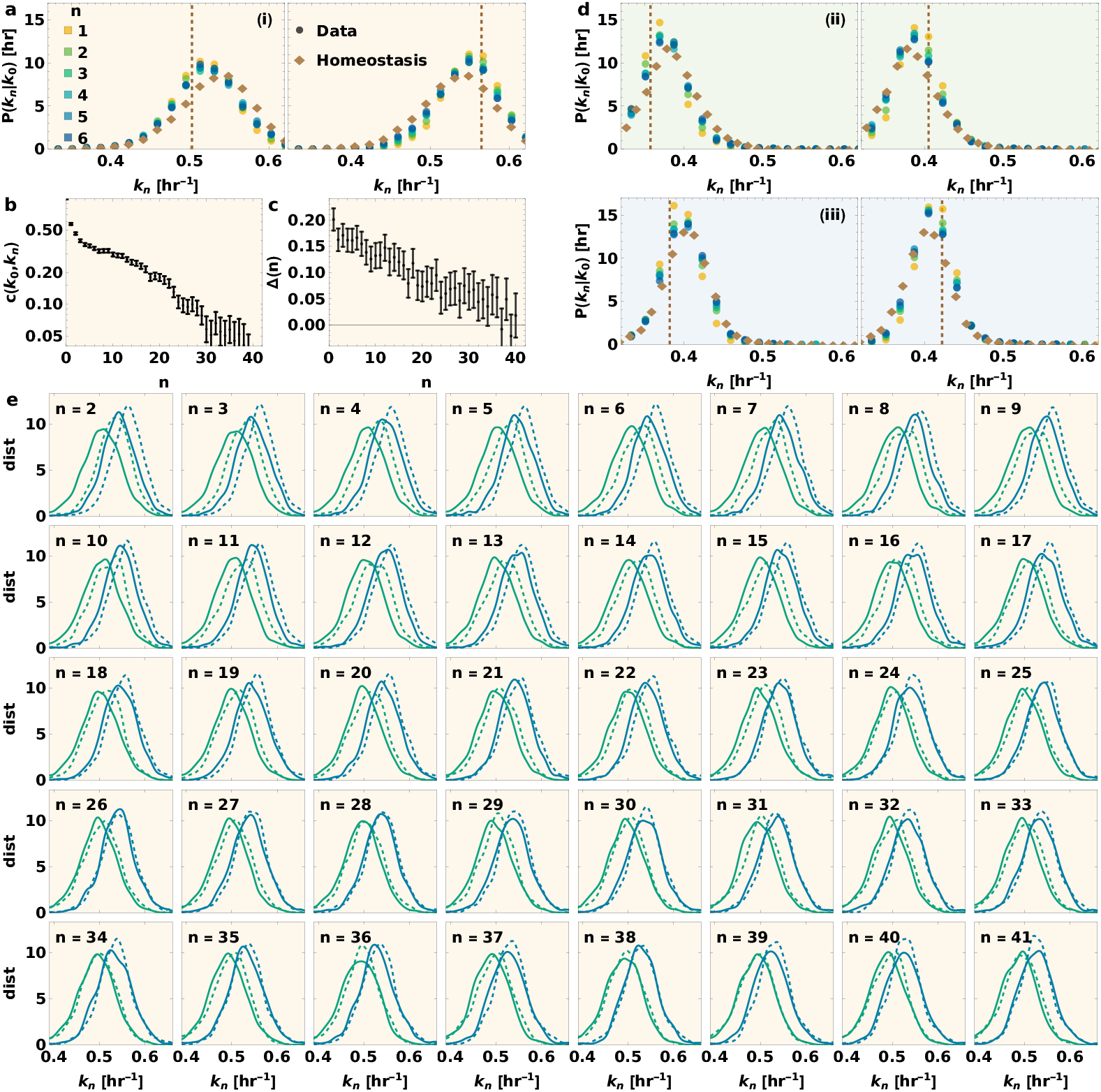
Stochastic intergenerational homeostasis of the individual cell’s growth rate is sustained via the plastic, reflective, memoryful scheme. (a) The intergenerational evolution of the distribution of growth rates, k_n_, starting from a given growth rate in the starting generation (location of dashed line), for cells in complex media at 30° C. n denotes the generation number, with the starting generation labeled by n = 0. The growth rate distributions do not converge to the steady state distribution (diamond points) even after 6 generations. The growth rate in the starting generation is the first quartile for the panel on the left, and the third quartile for the panel on the right. (b) Coefficient of correlation between the growth rates in the initial (zeroth) generation and after n consecutive generations for different n values, plotted on a log scale. (c) Δ(n), a measure of the non-Markovian behavior of single cell growth rate given by (2), is plotted as a function of generation n, for cells in steady state. Δ(n) is zero for all n for Markovian systems, and a non-zero value for Δ(n) indicates the presence of non-Markovian memory up till generation n. (d) The same distributions as (a) for cells in (ii) dilute complex media and (iii) minimal media. (e) For cells growing in complex media in steady state, the conditional distributions of the n^th^ generation’s growth rates (k_n_) are plotted, conditional on whether the previous generation’s growth rate (k_n−1_) is greater than (blue) or less than (green) the median growth rate, and whether the initial generation’s growth rate (k_0_) is greater than (dashed) or less than (solid) the median. For a completely Markovian process, the distribution of k_n_ conditioned on k_n−1_ is completely independent of the growth rates of all generations prior to n −1, including k_0_. Thus the difference between solid and dashed curves indicates the presence of inter-generational memory in growth rate. Intergenerational growth rate follows nonMarkovian dynamics, retaining memory up to ∼ 40 generations.

### A metric for characterizing the degree of non-Markovianity

We introduce and define the following measure:

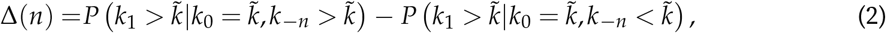

where, 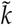 is the population-wide median single cell growth rate. Starting from the (arbitrarily chosen) zeroth generation, *k*_*i*_ is the growth rate of the *i*^th^ generation. Here, positive *i*’s correspond to generations following the zeroth generation, while negative *i*’s correspond to generations prior to the zeroth generation. We note that the choice of 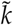 here is not unique, and could potentially be replaced with any other constant value. We choose the population-wide median for this value to ensure that we have sufficient data when selecting points with 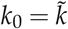. For a completely Markovian intergenerational process the probability distribution of *k*_1_ depends only on *k*_0_ and is independent of *k* _−*n*_. Thus Δ(*n*) is zero for all positive integers *n* for a Markovian process. Fig. 3 c shows that Δ takes around 40 generations to go to zero, indicating that single cell growth rate retains non-Markovian memory for 2 to 40 generations.

### Distinct signatures of long- and short-term memory retention in the individual cell growth rate

The generation-generation autocorrelation function of the individual cell growth rate displays a clear biphasic behavior (Fig. 3 b), suggesting that distinct underlying biological mechanisms are responsible for short-term (working) memory and long-term (fossilized) memory. There is a prominent “knee” separating the two regimes. Two putative power-law regimes are revealed in the log-log plot (Fig. 3 b). In contrast, the generation-generation autocorrelation dies out within ∼5 generations for the initial size dynamics (Fig. 2 d), matching the predictions of our theoretical framework which incorporates the assumption of Markovianity.

### Emergent simplicity: Intergenerational scaling law yields stochastic cell size homeostasis

We find that the conditional distribution of the next generation’s initial size, conditioned on the current generation’s initial size, undergoes a scaling collapse upon rescaling by the conditional mean:

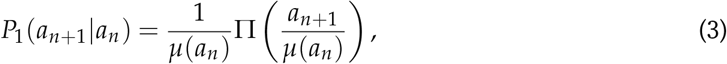

where Π denotes the new mean-rescaled distribution (with unit mean) and *µ*(*a*_*n*_) = ⟨*a*_*n*+1_ |*a*_*n*_⟩ the conditional mean of the initial size in the next generation, given the initial size in the current generation. We see direct evidence of this intergenerational scaling of initial sizes (Fig. 2 a– b). We thus have stochastic map for intergenerational cell size statistics of an individual cell in a given balanced growth condition, using as inputs the universal mean-rescaled distribution function, Π, and the calibration curve corresponding to the conditional mean, *µ*, for that growth condition (Mathematical Derivations section 3 and [31]). Since the conditional mean is found to have a linear form: *µ*(*a*) = *αa* + *β*, the stochastic map can be simply written down as:

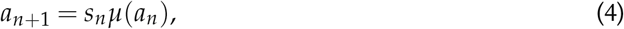

where the *s*_*n*_’s are independent identically distributed stochastic variables drawn from the meanrescaled distribution Π. We note that the shape of Π can vary from balanced growth condition to condition. (We discuss this fact further in a subsequent section on tradeoffs in stochastic intergenerational cell size homeostasis.) The significant advantage derived from the extraction of the intergenerational scaling law from data is that we are able to decompose a twodimensional function (the probability distribution of next generation’s initial size given the current generation’s initial size) into two one-dimensional functions – (i) the mean-rescaled distribution, Π, which is invariant with respect to the current generation’s initial size in a given balanced growth condition, and (ii) the linear calibration curve *µ*(*a*). The reduction in complexity achieved through this dimensional decomposition proves essential in obtaining further insights and practical predictions from our theory, since both the one-dimensional functions are readily estimable from data (Fig. 2 b, c). In contrast, estimating the full conditional distributions of next generation’s initial size given current generation’s initial size for every possible current generation’s initial size is practically unfeasible even for datasets from high precision high throughput experiments such as the ones we have presented here. Our fitting-free predictions match our data remarkably well for all conditions (Fig. 2 e).

### Stochastic homeostasis of the single cell growth rate

We can also provide direct evidence for stochastic intergenerational homeostasis of the single cell growth rate in the same datasets using analysis similar to what we previously presented for the case of cell size homeostasis (Fig. 1 h, i, j, l, m). But unlike cell size, which sustains homeostasis through a Markovian process (Fig. 1 k), the dynamics of growth rate are non-Markovian and retain memory for up to ∼40 generations (Fig. 3). This difference is evident in the generational progression of cell size and growth rate in reaching the steady state distribution starting from different initial conditions. While cell size attains homeostasis in ∼6 generations for all growth conditions (Fig. 2 e), the growth rate distributions take several times this generational timescale to reach the steady state distribution (Fig. 3 a, d). The theoretical formulation we previously presented to predict cell size dynamics across generations, if applied to the growth rate dynamics, would predict a much faster convergence than observed. Moreover, the intergenerational scaling law governing cell size homeostasis is found to not be applicable as written to growth rate homeostasis. Thus a new theoretical formulation that also accounts for non-Markovian memory remains to be developed for cell growth rate homeostasis.

## Discussion and concluding remarks

### Stochasticity specifies the conditions for cell size homeostasis

The necessary and sufficient condition for stable cell size homeostasis is |*α*| ≤ 1/*s*_*max*_ (see [31] for derivation), where *α* is the slope of the linear conditional mean, *µ*(*a*) = *αa* + *β*, and *s*_*max*_ is the upper bound of the mean rescaled distribution Π. This condition ensures that all moments of the steady state initial size distribution converge to finite values independent of the starting initial size. This is a stronger bound on *α* than that obtained from the quasi-deterministic framework, *α <* 1 [26, 32–34], which only ensures the convergence of the mean in our framework.

### Tradeoffs in stochastic intergenerational homeostasis

Although our framework predicts an upper bound for |*α*| for cell size to be in homeostasis, there is no lower bound. The smaller the *α*, the faster the convergence to homeostasis, and for *α* = 0, the system converges to the homeostatic distribution within one generation starting from any arbitrary initial condition. Yet, under all growth conditions, we observe that *α* always lies remarkably close to our theoretically predicted upper bound (Fig. 2 f), suggesting that speeding up the convergence to homeostasis comes at a cost to the cells, resulting in the slowest convergence possible while also maintaining stable homeostasis: a precision-energy tradeoff. The upper bound 1/*s*_*max*_ is inversely related to the degree of noise in the mean rescaled distribution Π. Thus more noise in this distribution enforces a faster convergence to homeostasis.

Another tradeoff we extract from data is the positive correlation between the coefficient of variation of the mean rescaled distribution Π (which is a measure of the noise in the homeostatic control of cell size) and the average single-cell growth rate for cells growing in different balanced growth conditions (Fig. 4). Faster growth comes at the expense of increased noise: a precisionspeed tradeoff [35].

**Figure 4:**
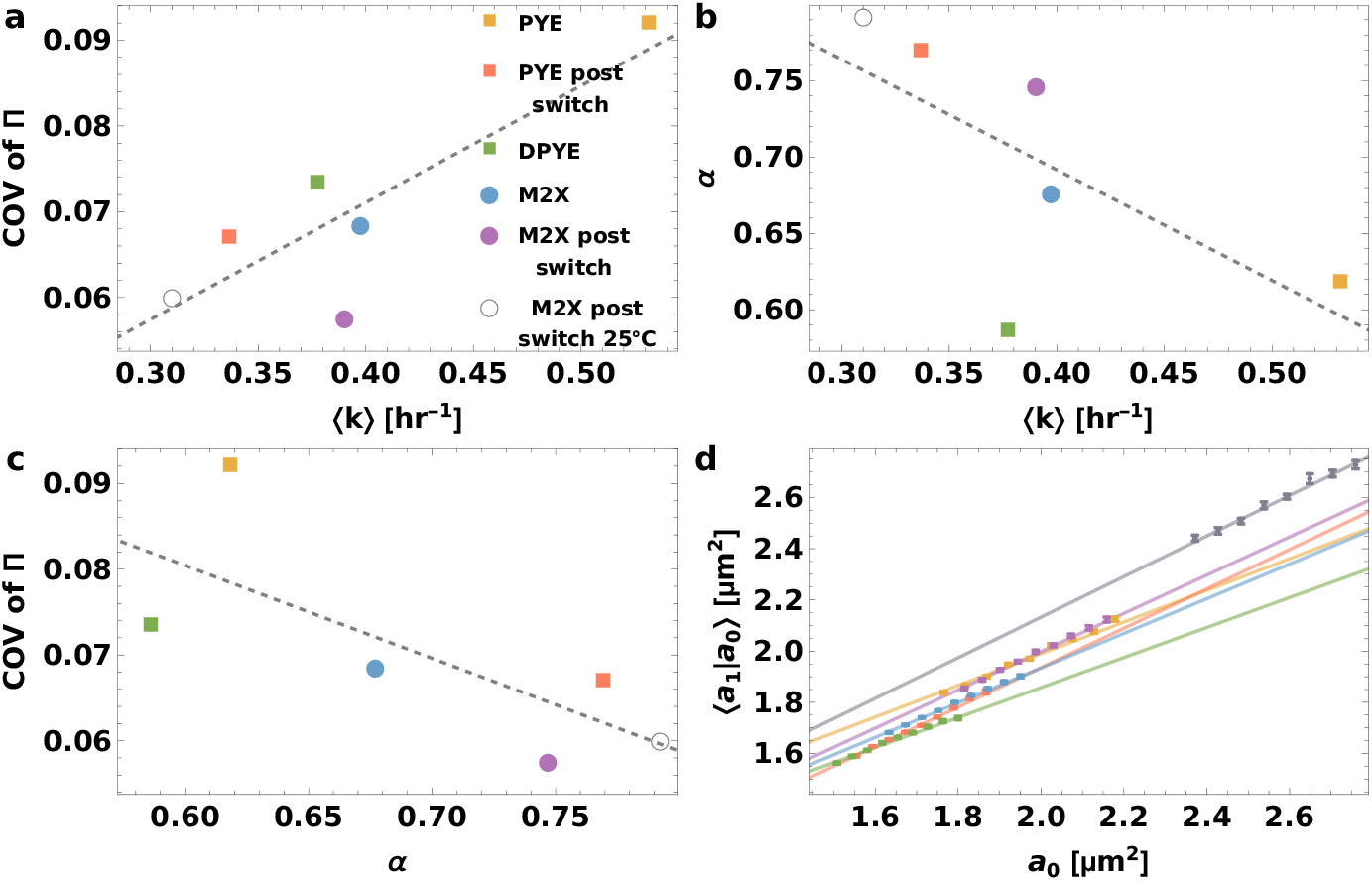
Observed tradeoffs in stochastic intergenerational cell size homeostasis. (a) The coefficient of variation of the mean rescaled distribution Π which determines the noise in the steady state initial size distribution is plotted as a function of mean growth rate (averaged over the entire population) for different growth conditions. Here, PYE represents complex media, represented by squares, and M2X represents minimal media, represented by circles. D represents dilute, filled points represent growth conditions at 30° C, and the empty point 25° C. The dashed line shows the linear fit. The positive slope indicates COV increases as growth rate increases. (b) α, which is the slope of the linear fit of mean subsequent generation’s mean rescaled initial size (a_1_), conditioned on the current generation’s mean rescaled initial size (a_0_), as shown in (d), is plotted as a function of mean growth rate. α determines the number of generations to reach steady state. The decrease in α with increasing growth rate indicates that it takes more generations to reach steady state when growth is faster. (c) COV is plotted as a function of α. (d) The binned means of the subsequent generation’s initial size are plotted as functions of the current generation’s initial size (background scatter), with error bars showing standard error, along with the linear fit. The different colors correspond to the different growth conditions in (a).

### Broad applicability and mechanistic basis of the intergenerational scaling law governing stochastic cell size homeostasis

The intergenerational scaling law governing cell size homeostasis is found to be broadly applicable across growth conditions (Fig. 2), bacterial species [31], and microenvironments [36]. Thus it provides a universal solution to the problem of stochastic intergenerational bacterial cell size homeostasis despite the differences in the specifics of the underlying molecular circuitry across different bacteria and even across different growth conditions for the same bacterial species. These emergent simplicities arise due to the fact that the organizational motif representing the nature of coupling of growth to division is effectively the same in all of these scenarios, despite apparent differences in actualization through molecular circuitry in each case. As we have demonstrated in [18], systems in which division is triggered through a first passage process of a protein crossing a fixed threshold amount followed by a constant time delay exhibit the same scaling laws demonstrated here. The identity of the protein may vary across organisms and growth conditions, with different accumulation rates and different time delays post crossing the threshold, which result in the differences between the mean rescaled distributions Π and the linear conditional mean *µ*(*a*) across these conditions. A potential candidate that exhibits the properties required by the thresholding protein in *C. crescentus* cells is FtsZ [18].

## Author contributions

S.I.-B. conceived of the research; K.J., R.R.B. and S.I.-B. designed the research; K.J. extracted the intergenerational scaling law from the data and performed data analysis under the guidance of S.I.-B.; K.J., R.R.B. and S.I.-B. developed the theoretical framework and performed analytic calculations; C.S.W. and S.I-.B. designed the experimental setup; K.F.Z. and C.S.W. actualized and refined the experimental pipeline under the guidance of S.I.-B.; C.S.W. and S.E. designed and developed the analysis pipeline with input from R.R.B. and S.I.-B.; K.F.Z., C.S.W, E.S., J.C. and S.I.-B. performed experiments; K.J., K.F.Z., S.R., R.G. and J.S. contributed to manual supervision of image analysis; K.J., C.S.W., R.R.B. and S.I.-B. wrote the paper with input from all authors; S.I.-B. helped shape the research and analysis and supervised all aspects of the research, and saw it through to fruition.

## Acknowledgements

We are grateful to the Iyer-Biswas group members for insightful discussions and feedback. S.I.-B. thanks Daniel S. Fisher for useful discussions. We thank Purdue University Startup funds, Purdue Research Foundation, the Purdue College of Science Dean’s Special Fund, and the Showalter Trust for financial support. K.J. and S.I.-B.acknowledge support from the Ross-Lynn Fellowship award. S.I.-B. thanks the Purdue Physics and Astronomy REU program for hosting and providing financial support for R.G. and J.S’s contributions. S.E. and S.I.-B thank Purdue’s Data Mine Learning Community for computational resources.

## Supplementary Information

### Mathematical Derivations

#### 1. Notation

We use *a*_*i*_ to denote generic size at birth (initial size) and *a* _*f*_ to denote size at division (final size); *a*_*n*_ (where *n* is distinct from *i* or *f*) is a random number that denotes the initial size in the *n*^th^ generation; 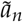 is a random number that denotes the final size at the end of the *n*^th^ generation; *r* is a random number that is the division ratio, the ratio of the initial size to the final size in the previous generation, defined using the relation 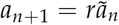; *P*(*X*) is the usual notation denoting the probability distribution of random variable *X*; *P*(*X* | *Y*) denotes the conditional probability distribution of *X* conditioned on *Y*; ⟨*X*|*Y*⟩ denotes the conditional average of *X* conditioned on *Y*; *P*_*f*_ (*a*_*f*_| *a*_*i*_) denotes the probability distribution of *a* _*f*_ conditioned on *a*_*i*_ within the same generation; *P*_*n*_(*a* |*a*_0_) is the probability distribution of *a*_*n*_ at *a* conditioned on *a*_0_, the initial size in generation 0, often referred to as the (arbitrarily chosen) “initial” generation; and *P*_*r*_(*r*) denotes the probability distribution of the division ratio *r* in any generation, while *P*_*r*_(*r*|*a* _*f*_) is the distribution of *r* conditioned on the corresponding final size *a* _*f*_.

#### 2. Derivation of intergenerational framework

Since in balanced growth conditions the sequence of initial cell sizes *a*_0_→ *a*_1_→ … →*a*_*n*_ occurs through a Markov process (i.e., is a Markov chain), the probability distribution of the final generation size, *a*_*n*_, can be expressed as a convolution of conditional probabilities governing each step in the sequence:

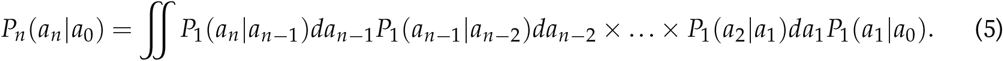

This is the same as Eq. (1) in the main text.

#### 3. Relating the functions Π and *µ* to Π _*f*_ and *µ*_*f*_

We begin with the emergent simplicity of mean rescaling, expressed for initial and final sizes in generation *n*:

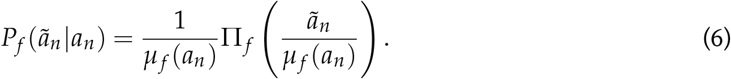

The initial size in the next generation, *a*_*n*+1_, can be found from the final size, 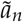, in generation *n*, using the division ratio 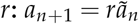. Given that *r* is a random variable with distribution *P*_*r*_(*r*) that is independent of 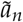, we can relate the mean of *a*_*n*+1_, *µ*(*a*_*n*_), to the mean of 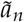, *µ*_*f*_ (*a*_*n*_), as follows:

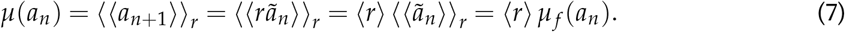

Note that there are two steps of averaging; the inner angular brackets average over possible growth-division processes at fixed *r*, whereas the outer angular brackets independently average over *r*. Meanwhile, the conditional probability distribution of *a*_*n*+1_ given *a*_*n*_ becomes:

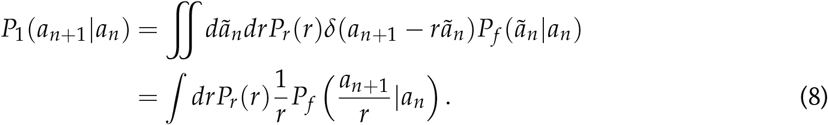

Herein, *δ* denotes the Dirac delta function. Using Eqs. (6) and Eq. (7),

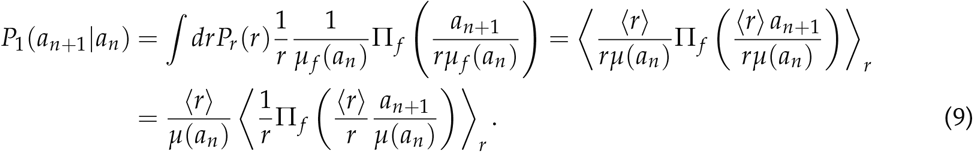

We have thus arrived at the final intergenerational scaling form in the main text, Eq. (3), with the scaling function Π(*s*) defined (setting *s* = *a*_*n*+1_/*µ*(*a*_*n*_) in the above) as:

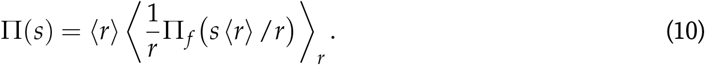

#### 4. Deriving the coefficient of correlation

Since the conditional mean of subsequent generation’s initial size given current generation’s initial size has the linear form *µ*(*a*) = *αa* + *β*, thus, for *n > m*,

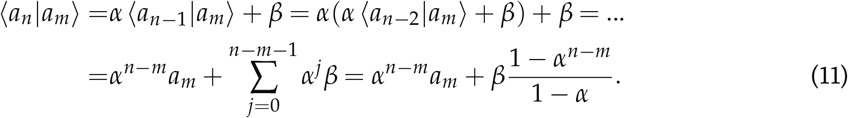

Now, the coefficient of correlation is,

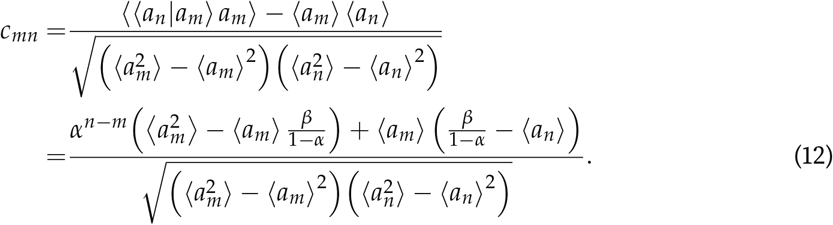

The distribution of initial sizes is invariant in steady state, thus 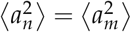 and ⟨*a*_*n*_⟩ = ⟨*a*_*m*_⟩ = *β*/(1 − *α*) (from the linear conditional mean ⟨*a*_*m*_⟩ = *α* ⟨*a*_*m*_⟩ + *β*). Thus,

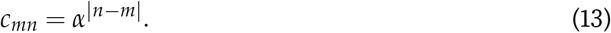

## Experimental Methods

The basic protocol for using a SChemostat device for collecting high throughput single-cell data under constant growth conditions is described in [15]. For datasets involving switches from one condition to another the following modifications were made to this basic protocol. First, a two input PDMS device was fabricated and the tubing was connected. Next, the channels were cleaned by connecting syringes to the outlet of the device and flushed with KoH, ethanol, and DI water, in order. The DI water syringe was left connected to the device and syringe pump was left running while connecting the input tubes to each other. The pumps were stopped right after the first tube was connected. Next, the PDMS device with connected DI water syringes was placed onto the microscope stage and the channel was brought in focus. Next, the cells were induced into the first channel through the following steps in order: (i) the DI water syringe was disconnected from the needle tip and connected to a 1mL cell-culture syringe, (ii) the input tubes were disconnected from each other and the cell culture medium was flushed from the outlet side through the channel, and (iii) the input tubes were re-connected after 0.75 mL of media was pushed into the channel. These steps were then repeated for all the other channels. Next, after waiting for 1h until the desired cell density was reached (which was checked occasionally via eye port), the prepared media syringes were set up in the syringe pump and the airless tubing was connected to each other. Finally, the media syringes were connected to one channel at a time through the following steps in order: (i) all media syringe tubing were disconnected from each other and set to a flow rate of 50L/min, (ii) after waiting for the media to flow out of the tubing from every syringe, the re-connected input tubes of one channel were disconnected, (iii) the plunger of a cell culture syringe was slowly pushed until droplets formed on the input tubing, (iv) one of the media syringe tubes was connected to one of the input tubes of the channel, (v) upon confirming that media was released at the opposite input tubing and waiting for 1min of media flow, the second media syringe tube was connected to the other, still open input tube, (vi) the cell-culture syringe and needle tip was removed immediately after from the output side of the device, then (v) it was confirmed that media from both syringes flowed through the channel and was released at the output side. These steps were repeated for the other channels. Once the media flowed through all desired channels, the syringe pump containing the second media was stopped and the cells were subjected to only the first media. Then the flow conditions were set up and the experiment was started.

